# Cyclophosphamide induces the loss of taste bud innervation in mice

**DOI:** 10.1101/2023.06.14.544942

**Authors:** Ryan M. Wood, Erin L. Vasquez, Eduardo Gutierrez Kuri, Kevin Connelly, Saima Humayun, Lindsey J. Macpherson

## Abstract

Many common chemotherapeutics produce disruptions in the sense of taste which can lead to loss of appetite, nutritional imbalance, and reduced quality of life, especially if taste loss persists after treatment ends. Cyclophosphamide (CYP), an alkylating chemotherapeutic agent, affects taste sensitivity through its cytotoxic effects on mature taster receptor cells (TRCs) and on taste progenitor cell populations, retarding the capacity to replace TRCs. Mechanistic studies have focused primarily on taste cells, however, taste signaling requires communication between TRCs and the gustatory nerve fibers which innervate them. Here, we evaluate the effect of CYP on the peripheral gustatory nerve fibers that innervate the taste bud. Following histological analysis of tongue tissues, we find that CYP reduces innervation within the fungiform and circumvallate taste buds within 4-8 days after administration. To better understand the dynamics of the denervation process, we used 2-photon intravital imaging to observe the peripheral gustatory neuron arbors within individual fungiform taste buds before and after CYP treatment. We find that gustatory fibers retract from the taste bud proper but are maintained within the central papilla core. These data indicate that, in addition to TRCs, gustatory nerve fibers are also affected by CYP treatment. Because the connectivity between TRCs and gustatory neurons must be re-established for proper function, gustatory fibers should continue to be included in future studies to understand the mechanisms leading to chemotherapy-induced persistent taste loss.

## Introduction

50-80% of cancer patients who receive chemotherapy treatments experience taste alteration in the form of reduced taste sensitivity, distorted taste perception, or even complete loss of taste during treatment^1-4^. These deficits can persist for months beyond the end of therapy^4^. Indeed, the side effects of chemotherapy on taste are listed as one of the most disturbing elements of treatment, negatively impacting the patient’s ability to eat and drink and maintain healthy nutritional balance^5^. This can result in loss of appetite, malnutrition, poor recovery, and a reduced quality of life. Unfortunately, there are few, if any, effective treatments for those experiencing taste loss, partially due to the major gaps in the understanding of the mechanisms leading to persistent taste dysfunction after cancer treatment.

Since many chemotherapy drugs target the fast dividing cancer cells, other proliferative cell types are also affected, most visibly causing hair loss, but also affecting the immune system, digestive system, and the highly proliferative taste progenitor cells within taste buds^6^. Alkylating chemotherapy drugs such as cyclophosphamide (CYP) produce a two-phase disturbance in taste behavior in mice – one immediately following drug administration, and a second which emerges several days later ^4,6-9^. The first phase is due to direct cytotoxic effects on the lingual epithelium causing an early loss of taste receptor cells. Then, the second disturbance is due to the disruption of the taste stem cell population responsible for TRC renewal, temporarily retarding the system’s capacity to replace the TRCs^4,7-9^. Once chemotherapy treatment ceases, the remaining population of unaffected taste stem cells can begin to repopulate the taste buds with new TRCs. The normal rate of turnover for TRCs in the taste bud is 10-20 days^10^. In mouse models of CYP taste loss, most TRC populations recover 16-20 days after treatment^7-9^.

Another type of chemotherapy drug, used most often to treat basal cell carcinomas, inhibits the hedgehog signaling pathway. Notably, these drugs, including vismodegib and sonidegib, produce a significant disturbance or complete loss of taste in most patients within 2 months after the initiation of treatment^11-13^. In this case, these drugs do not produce the initial die-off of TRCs but cause a reduction in the proliferation of taste progenitor cells and delay TRC maturation^14^. Regeneration of the TRCs after treatment depends on Sonic Hedgehog (Shh) signaling supplied by the gustatory neuron fibers^14^. A careful analysis of chorda tympani fibers within fungiform taste buds after hedgehog pathway inhibition with sonidegib showed a significant effect on the innervation pattern within the papillae after the loss of TRCs. After the loss of the TRCs, the fibers project outward into a flat disc-shaped bundle that doubled in size^15^. The authors suggest that maintenance of fibers near the sites of lost taste buds is important for regeneration, such that the continued presence of these fibers in the vicinity of regenerating taste buds promotes reinnervation of newly formed buds.

In any case, as TRCs repopulate within the taste buds, they must resume communication to peripheral gustatory sensory neurons innervating the tongue via synaptic contacts. If this rewiring process is successful, taste function will gradually return to baseline. However, the fact that some chemotherapy patients experience protracted taste disturbances (over months and years) indicates that this recovery and rewiring process is prone to failure^5^. To date, most research on mechanisms leading to chemotherapy-induced taste dysfunction has focused on the loss and repopulation of TRCs within taste buds. A critical element that is missing in these studies is understanding the contribution of gustatory sensory neurons in the initial loss and eventual re-establishment of taste function after chemotherapy treatment.

To monitor the morphology of gustatory nerve fibers innervating the taste buds and to determine if the innervation patterns change over the course of CYP treatment, we used a combination of immunohistochemistry and intravital 2-photon imaging. We hypothesized that as taste receptor cells die after CYP treatment, gustatory fibers would retract from the taste buds, but would eventually re-innervate the buds as new taste receptor cells matured and repopulated the buds. Our results indicate that this hypothesis, where we see that as the taste receptor cells die off, the gustatory fibers innervating them are lost. However, both taste buds and the gustatory fibers are variability in how they are affected by treatment, as some maintain their cells and innervation and others are completely lost. This needs to be followed up on, to see how these taste cells recover and if they can regain innervation.

## Materials and Methods

### Animals

All procedures were conducted in accordance with US National Institutes of Health (NIH) guidelines for the care and use of laboratory animals and were approved by the University of Texas at San Antonio IACUC. Both male and female mice were used in this study. For immunohistochemical experiments of sectioned tongue tissues, C57/b6 mice (Jax Strain #000664) were used along with Phox2b-Cre; Ai9 double transgenic mice (Jax Strains #016223, #007909). All mice were between 3–8 months old; weighing between 18-35g. Food and water were available ad libitum additionally after injection they were given access to a nutrient supplement (DietGel®) to aid recovery. The mouse colony was maintained on a regular 12/12 hr light-dark cycle.

### Drug treatment

Cyclophosphamide solution was prepared immediately prior to a single IP injection by dissolving cyclophosphamide monohydrate powder (Cat. No. AC203960250, Fisher Scientific, Hampton, NH, USA)^7-9^ in sterile saline (0.9%) for a dose of 100mg/kg. Day 0 Control group for tubulin experiments were not injected. For Phox2B-Ai9 mice, control mice were injected with sterile saline based on their weight to match cyclophosphamide injected mice. Mice were observed each day after injections through weighting and physical wellbeing checks.

### Sample collection

Mice were euthanized through CO2 inhalation followed by trans-cardial perfusion with 0.1 M PBS and 4% paraformaldehyde solution in 0.1 M PBS, sequentially. Tongues were extracted and soaked in 4% paraformaldehyde solution for 2 hours at room temperature then cryoprotected in 30% sucrose at 4C overnight. The tongues were then dissected to separate the circumvallate papillae (CV) and the anterior 2/3 of the tongue housing the fungiform papillae. The anterior 2/3 of the tongue was then cut sagittal to split it into left and right halves. The left and right tongue tip and the CV were embedded in OCT compound and stored at –80° C. Samples were cryo-sectioned at 15μm thickness and mounted directly on slides.

### Immunostaining

Sections of the CV and tongue tip were blocked in 10% donkey serum at 4°C for 1 hour before they were incubated with primary antibodies: TRPM5 (1:500; provided by Zuker Lab^16^), Beta Tubulin III (1:200; Abcam ab41489) and Car4 (1: 1000; R&D Systems AF2414), or with Troma (1:1000; DSHB Cat# TROMA-I, RRID:AB_531826) in 10% donkey serum in 0.1 M PBS overnight at 4ºC. After 3x 5-minute washes with PBS, secondary antibodies (Alexa Fluor®chicken488 AB2340375, Alexa Fluor® goat594 AB2340432, Alexa Fluor® rabbit647 AB2492288) were added (1:1000) in 10% donkey serum in PBS and incubated overnight at 4°C. The slides were then washed 3x 5 minutes with PBS and then mounted for imaging. A Zeiss 710 confocal microscope was used to image the tongue sections. Z-stack images consisted of 15 stacks at 1um depth.

### Whole mount immunofluorescence

The right tongue tips were thinned to the dorsal epithelial surface through removal of the ventral fat. The Epithelium was placed in a 24 well plate and incubated with primary antibodies for Troma (1:500; DSHB Cat# TROMA-I, RRID:AB_531826) in 10% donkey serum in 0.1 M PBS for 4 days at 4ºC. After washing Tissues 3x with PBS for 20 minutes, secondary antibodies (Alexa Fluor®rat488; Thermo Fisher Scientific, catalog # A-11006, RRID AB_2534074.) were added (1:1000) in 10% donkey serum and incubated for 3 days at 4°C. The tissue was then washed 3x with PBS for 20 minutes and mounted on slides. A Zeiss 710 confocal microscope was used to image the tongue whole mounts. Z-stacks consisted of 50 stacks∼ at 1um depth, covering the tip of the taste bud to gustatory nerve fiber core.

### Immunohistology cell counting and quantification

For image analysis ImageJ (FIJI) was used. First, the number of TRPM5, CAR4 or TROMA positive cells in each taste bud section were counted. 20 buds per mouse were averaged for circumvallate tissues, and 10 taste buds were averaged for fungiform taste buds. To analyze the gustatory nerve fibers (either stained with tubulin or had tdTomato expression within the Phox2b cell population), each image stack was converted to a maximum projection image. The fluorescence was adjusted to a median brightness value and a threshold was set to median brightness. An ROI of each bud was measured and converted from pixels to um giving us a fractional area of the gustatory fibers, allowing quantification of the total gustatory nerve fibers in each bud. For tubulin experiments, days 4, 8, and 20 after cyclophosphamide injection were compared to uninjected controls using a Kruskal-Wallis test with Post-Hoc Dunn’s multiple comparison test. For Phox2b-Ai9 immunohistology experiments, cyclophosphamide injected mice were compared to saline injected controls using Mann-Whitney test. For Two-photon Experiments, individual taste buds were compared on Day 0 and Day 4 using Wilcoxon test. All tests were done using Graphpad Prism Software.

### Tongue imaging apparatus

A custom 3-D printed apparatus to stabilize the head and extrude the tongue was created with tinkerCAD software. The tongue holder is 2cm by 4cm and the hole for the tongue is a few millimeters in diameter. There are two additional holes for 2 metal screws to position and secure the mouse head. Magnets are attached to the screws and metal stage to stabilize the apparatus. The coverslip keeps the tongue flat to promote 3-D reconstruction of the gustatory fibers with the 2-photon microscope.

### Anesthesia and preparation of the mouse for imaging

Mice were given an initial IP injection of 10mg/kg ketamine and 0.1mg/kg xylazine (Covetrus). A IP injection was also given prior to positioning the mouse in the imaging apparatus to maintain fluid levels during imaging. The mouse was placed on his/her back and the tongue was inserted in the opening of the tongue holder using blunt forceps. After the tongue was placed through the opening, the coverslip was placed on the surface of the tongue. Initially, a 10x Olympus lens was used to spatially recognize that the same taste bud was being imaged. Once the correct bud was identified, a 40x water immersion lens was used to image the gustatory fibers. After each imaging session, the mouse was given an IP injection of Atipamezole (Covetrus) (1mg/kg) to reverse the effects of the sedative.

### 2-photon imaging

2-photon excitation microscopy imaging was used to obtain 3-D images of a live mouse tongue, a Bruker system with prairie view software. A laser strength of approximately 400mV and a wavelength of 1100 nm was used to visualize the tdTomato labeled nerve fibers innervating the tongue. The same PMT and imaging strength was used each day to ensure no variation due to the setting changes. Images were collected at 1024x1024 pixels, resonance galvo, with resonance averaging at 2 frames. This resonance setting was used because it takes a faster image which is critical due to our time constraints of the sedative. Z stacks were collected at .5um intervals covering approximately a volume of 100um, cubed.

Typically, most of the taste bud is imaged within the first 50um of the Z-axis, but additional images were collected toward the base of the bud to help orient the 3-d volume as well as confirm that the same bud had been imaged day to day.

### 3-D image analysis

ImageJ and Imaris were utilized for data analysis and 3-D reconstruction of whole-mount and 2-photon images. ImageJ software was used to obtain a volume measurement of the fibers in pixels, cubed. First, a consistent threshold setting was used in ImageJ to minimize background fluorescence/autofluorescence. Then, ImageJ was used to extract the region of interest, and remove extraneous pieces that were not included in the analysis. This region of interest excluded the bottom portion of the bud that comprises the dense papilla core. In addition, the extragemmal fibers that surround the bud were also excluded. A custom volume calculator macro in ImageJ was utilized to calculate the volume of fibers innervating each taste bud. Imaris software was used to create 3-D reconstructions of the innervating fibers, after ImageJ was used to select the region of interest. The “surface volume” setting was applied to the z-stack. This setting finds the edges of 3-D images and renders an outline of the 3-D image.

## Results

### Cyclophosphamide Treatment Causes a Transient Loss of TRCs and Sensory Fiber Innervation in Circumvallate Taste Buds

It is well established that a single treatment of cyclophosphamide causes a transient loss of taste receptor cells within lingual taste buds^17^. Slight differences in the timing have been reported for Type II and Type III TRCs^9^. To determine whether the gustatory neurons are also affected by CYP treatment, we used an approach similar to Mukherjee et al.^7-9^, injecting a single dose of CYP (100mg/kg, IP) and examining TRCs and neuronal fiber innervation at 0, 4, 8 and days post-treatment (Figure 1A). Like previous reports, we find that the average number of both Type II (labeled using an antibody against TRPM5), and Type III (labeled using an anti-Car4 antibody), are significantly reduced in circumvallate taste buds between 4-8 days after treatment, and cell numbers are re-established in the buds by 20 days after treatment (Figure 1C,1D; p<0.05).

**Figure 1:**
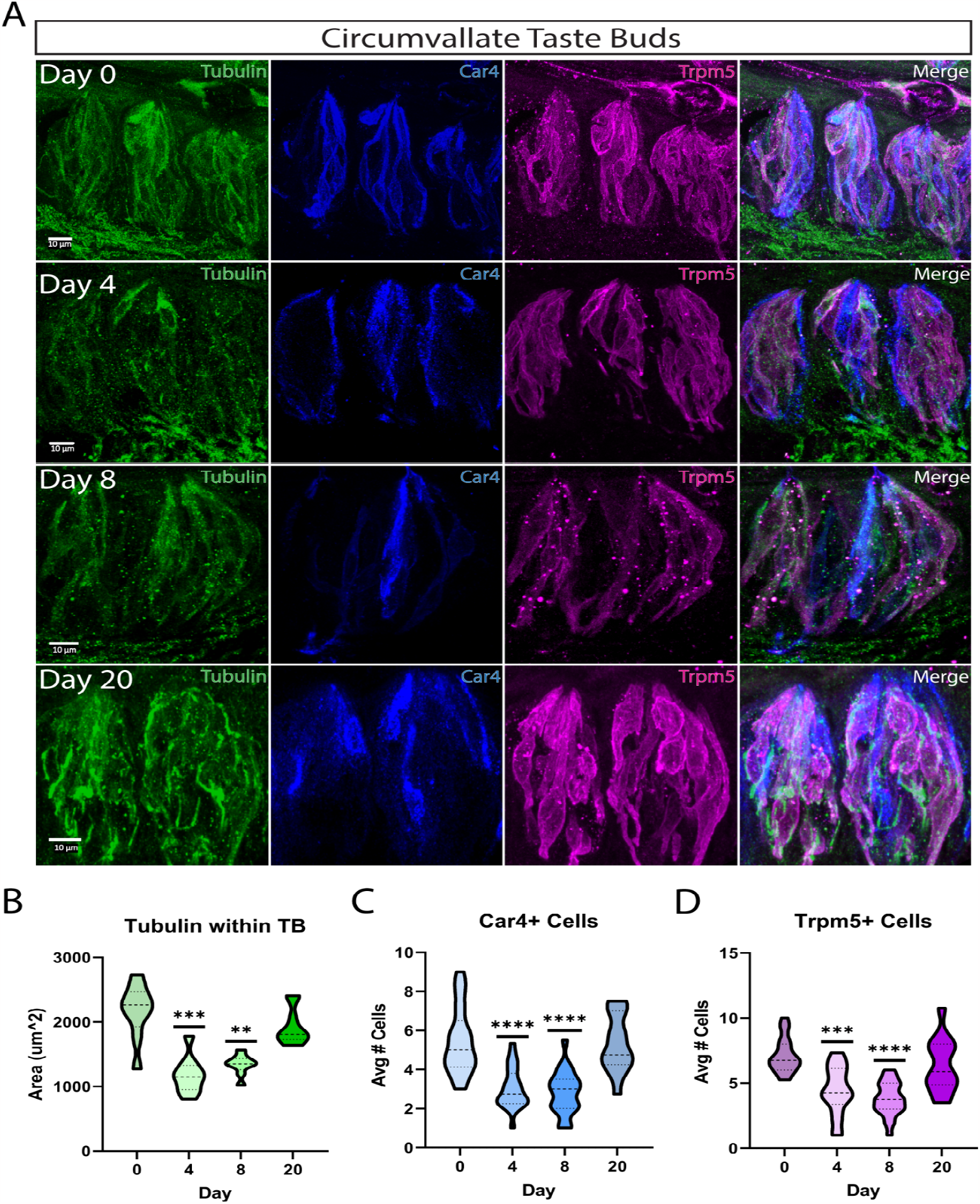
Cyclophosphamide Treatment Causes a Transient Loss of TRCs and Sensory Fiber Innervation in Circumvallate Taste Buds. **A**. Combined immune histology of circumvallate tongue sections stained with beta-tubulin (green) for neuronal fibers, Car4 (blue) for type III TRCs, Trpm5 (magenta) for type II TRCs. Sections were from C57 mice 0, 4, 8, and 20 days post-CYP IP injections (100 mg/kg). **B**. Violin plot quantification of the area of Tubulin within taste buds per day, (N=36 Taste Buds per group). **C, D**. Violin plot quantifications of the average number of TRC cells labeled for Car4 and Trpm5 within taste buds per day (N=78-86 Taste buds per group). Mann-Whitney Test were used to show significance for each graph (*p<0.05).

Using b-tubulin, a general antibody to label axons^18^, we find that the area labeled by b-tubulin within individual circumvallate taste buds is reduced in a similar fashion at 4 and 8 days after CYP treatment compared to day 0 controls (Figure 1B; Median D0: 2167.79 um^2^ D4: 1177.75 um^2^ D8: 1339.07 um^2^ D20: 1931.47 um^2^; p<0.05). It also appears that the peak of fiber reduction occurs at the day 4 time point, with the fibers’ area increasing slightly by day 8. By day 20, the area of innervation is not significantly different to controls. This indicates that peripheral sensory fibers are affected by CYP in a time course like that of the taste receptor cells.

### Phox2b-Ai9 Gustatory fibers are reduced in Fungiform and Circumvallate taste buds within 4 days of CYP Treatment

To further examine the reduction of gustatory nerve fibers affected by CYP treatment, Phox2b-Cre;Ai9 double transgenic mice were treated with CYP^19^. Phox2b-Ai9 Mice were injected with either cyclophosphamide or saline and sacrificed 4 days post-treatment (Figure 2A). Day 4 post-treatment was chosen as it was likely the peak of fiber loss.

**Figure 2:**
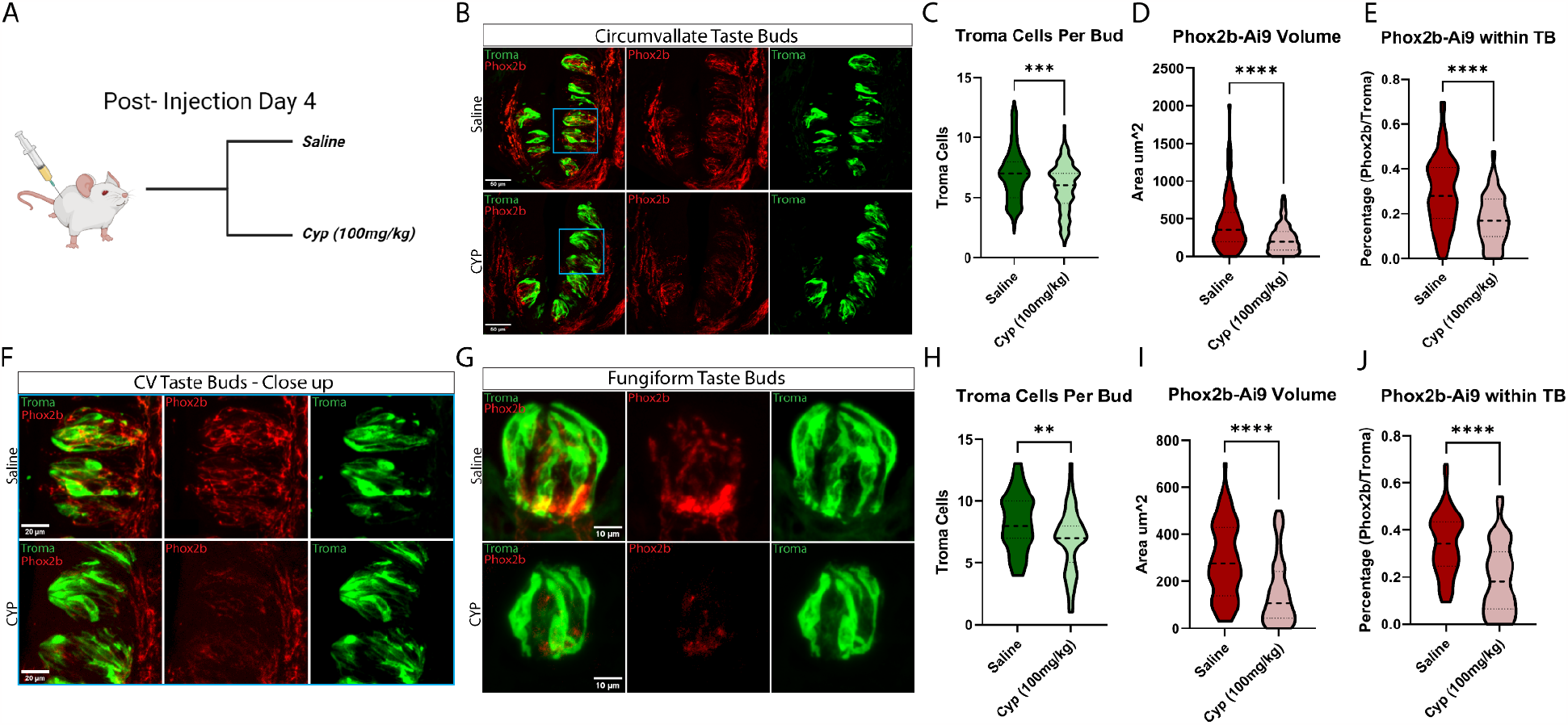
Phox2b-Ai9 Gustatory Fibers are Reduced in Fungiform and Circumvallate Taste Buds within 4 Days of CYP Treatment. **A**. Phox2b-Ai9 transgenic mice (Tdtomato expressed in Phox2b gustatory neuron population) were separated into two groups, injected with either Saline or CYP (100 mg/kg) (N=6 per group). **B**. Circumvallate tongue tections showing a single CV trench, stained with Troma (green) for taste receptor cells and Phox2b-Ai9 expression (Red) for gustatory fibers, 4 days post-injection. **C**. Violin plot of the average number of circumvallate TRCs labeled for Troma within taste buds for Saline vs CYP (N=111-113 TBs per group). **D**. Violin plot of the area of Phox2b within circumvallate taste buds for saline vs CYP, (N=111-113 TBs per group). **E**. Violin plot of the percentage of Phox2b area over the area of the circumvallate taste buds for saline vs CYP (N=111-113 TBs per group). Mann-Whitney Test were used to show significance (*p<0.05). **F**. Close up image of Circumvallate Taste Buds from part B, shown within the blue square. **G**. Fungiformtaste buds stained for Troma (green) for TRCs and Phox2b-Ai9 expression (red) for gustatory fibers from mice, 4 days post-injection. **H**. Violin plot of the average number of fungiform TRCs labeled with Troma for saline vs CYP(N=60 TBs per group). **I**. Violin plot of the area of Phox2b within fungiform taste buds for saline vs CYP (N=60 TBs per group). **J**. Violin plot of the percentage of Phox2b area over the area of the funfiform taste buds for saline vs CYP (N=60 TBs per group). Mann-Whitney Test were used to show significance for each graph (*p<0.05).

Circumvallate tissue was collected and analyzed from CYP and saline treatment groups (Figure 2B). We find that the average number of TRCs are reduced in the circumvallate taste buds by day 4 in our cyclophosphamide treatment group when compared to saline treatment mice (Figure 2C; Median CYP: 6 TRCs vs Saline: 7 TRCs; p<0.05). Examining the Phox2b-Ai9 signal, we find a reduction of the gustatory fibers innervating the taste bud 4 days after treatment in the CYP group (Figure 2D; Median CYP: 196.2 um^2^ vs Saline: 357.7 um^2^; p<0.05). Interestingly there is variability to this fiber loss when examining individual taste buds, as some of the taste buds retain gustatory fiber innervation, while others lost nearly all innervation (Figure 2D). This variability of nerve fiber loss among CYP treated animals also extends to TRCs loss. To quantify this, the percentage of gustatory fiber volume over the volume of the taste buds was used to normalize values. This resulted in a reduction of the percentage of fibers innervating taste buds (Figure 2B; Median CYP: 16.84% vs Saline: 28.05%; p<0.05). As before, TRC numbers decrease after CYP treatment. However, the gustatory fibers innervating the taste buds also decrease heavily afterwards within the circumvallate.

To see if CYP-induced fiber loss occurs in the anterior taste buds of the tongue, we examined the fungiform papillae (Figure 2G). Like the circumvallate taste buds, we found a decrease in TRC number within the fungiform taste buds (Figure 2H; Median CYP: 7 TRCs vs Saline: 8 TRCs; p<0.05). We examined the area of the gustatory fibers of the fungiform taste buds, also finding a reduction of the total fibers within the fungiform taste buds (Figure 2I; Median CYP: 106.8 um^2^ vs Saline: 275.3 um^2^; p<0.05), along with the percentage of fibers innervating the taste buds when compared to taste buds’ volume (Figure 2J; Median CYP: 17.94% vs Saline: 34.32%; p<0.05). The variability of taste bud/gustatory fiber loss found in the circumvallate taste buds is consistent in the fungiform taste buds as well. Interestingly, the number of empty cavities that once housed taste buds increased within the CYP treated mice, indicating a total reduction of taste buds induced by CYP treatment, as seen in a previous study ^7-9,20^. The observed loss in gustatory fibers innervating the fungiform papillae demonstrates that this phenomenon is not isolated in one region but appears to be affecting all the gustatory fibers within the tongue.

### Fungiform Taste Bud Density and Gustatory Fiber Volume Decreases Within 4 Days of CYP Treatment

Our initial experiments show a remarkable decrease in the gustatory fibers innervating fungiform taste buds within 4 days of CYP treatment. To understand the full scope of the CYP effects on gustatory neurons innervating fungiform taste buds, we performed a more in-depth analysis using whole-mount staining of the anterior tongue. Phox2b-Ai9 mice were injected with either CYP or saline, and the anterior tongue was collected 4 days post-treatment. An antibody for TROMA was used to visualize/analyze the taste receptor cells. Examining the 2mm tip of the anterior tongue, we found a decrease in the taste bud density 4 days post CYP treatment (Figure 3A,E; Median CYP: 4.635e-6 vs Saline: 6.612e-6; p<0.05).

**Figure 3:**
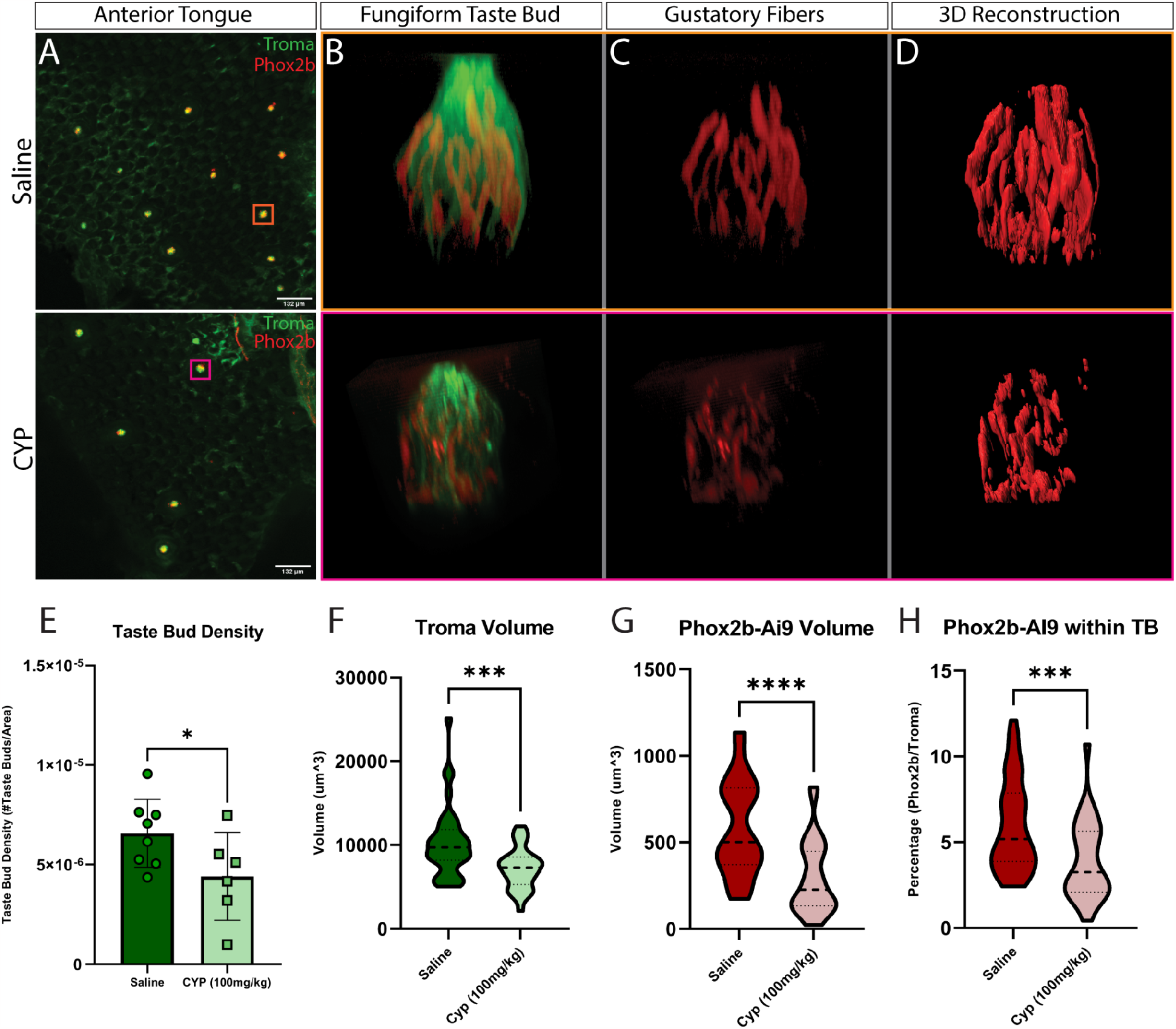
Fungiform Taste Bud Density and Gustatory Fiber Volume Decreases within 4 Days of CYP treatment. A-D.Top row: saline, bottom row: CYP. Tissues stained with Troma (green) to show TRCs and Phox2B-Ai9 expression (red) for gustatory fibers. **A**. Surface image of the anterior portion of a whole-mount tongue **B, C**. A single taste bud from each group showing gustatory fibers within the bud, along with showing an isolated reconstruction of only gustatory fibers. **D**. 3D reconstruction of a single taste gustatory fibers within taste buds. **E**. Density of taste buds within the first 2 mm of the anterior tip of the tongue for saline vs CYP (N=6-8 Mice per group). **F**. Violin plot of the volume of the taste buds for saline vs CYP (N=30 TBs per group). **G**. Violin plot of the volume of Phox2b-expressing fibers within taste buds for saline vs CYP (N= 30 TBs per group). **H**. Violin plot of the percentage of Phox2b-expressing fiber volume over the volume of the taste bud for saline vs CYP (N=30 TBs per group). Mann-Whitney Test were used to show significance (*p<0.05).

Whole taste buds were analyzed, finding a decrease in the total volume of the fungiform taste buds in CYP treated mice (Figure 3B, F; Median CYP: 9705 um^3^ vs Saline: 7325 um^3^; p<0.05). Similarly, the gustatory fibers were shown to decrease in volume after CYP treatment (Figure 3C,D,G; Median CYP: 226.5 um^3^ vs Saline: 501.3 um^3^ ; p<0.05). Normalizing to taste bud volume, the percentage of fibers innervating CYP treated taste buds also decreased when compared to saline controls (Figure 3H; Median CYP 3.255% vs Saline: 5.185%; p<0.05). This shows that CYP not only causes TRC death after treatment but induces the reduction of gustatory neuron fibers innervating taste buds to a similar or even greater degree.

### Using Intravital Imaging of Individual Fungiform Taste Buds to Visualize the Morphology and Measure The Volume of Gustatory Fibers Day to Day

Immunohistology is excellent for determining population changes from cyclophosphamide, however, it is also critical to determine how individual taste buds are affected by treatment over the course of multiple time points. To see the changes over time in individual taste buds, we used two-photon intravital imaging of the anterior tongue of anesthetized Phox2b-Ai9 mice with the use of a custom 3D printed head harness/tongue holder (Figure 4A). With this technique, we can track the same taste buds over the course of multiple days (Figure 4B). Z-stack images were taken from the tip of the gustatory fibers at the papillae pore opening, down the fibers’ core to examine both volume and morphology (Figure 4C). The volume of the innervating gustatory fibers remains consistent over the course of multiple days, fluctuating between 95%-105% of their original volume (Figure 4D). The volume of individual taste buds varies greatly, smaller taste buds having 1-2 individual fibers and larger ones having 20+ fibers making up their neuronal volume (Figure 4E). These results indicate that healthy taste buds remain relatively unchanged in their volume over time.

**Figure 4:**
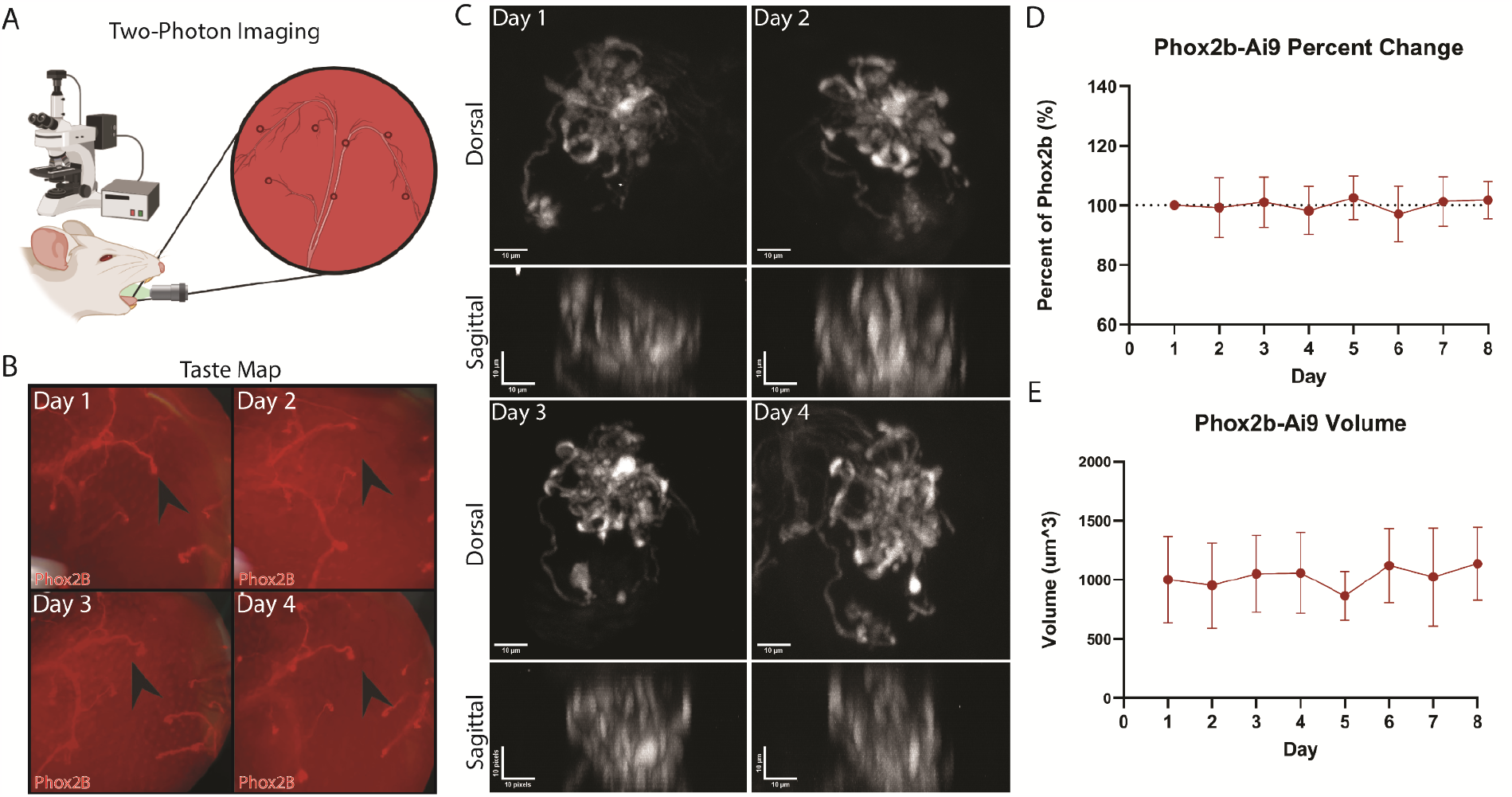
Using Intravital Imaging of Individual Fungiform Taste Buds to Visualize the Morphology and Measure the Volume of Gustatory Fibers Day to Day. **A**. Schematic of 2-photon imaging of the anterior fungiform taste buds from Phox2b-Ai9 mice **B**. Tongue map of a single mouse over the course of 4 days of imaging the anterior tongue. Nerve fibers are illuminated in red and single tracked taste buds are seen as bulbs indicated by the black arrow. **C**. A single taste bud tracked for 4 days. Dorsal view shows maximum projection of the z-stack. Sagital view shows maximum projection of a Z-stack. **D**. The percent change of taste buds over the course of 7 days, normalized to day 1 (N=12 Taste buds) along with standard deviation. **E**. The volume of taste buds over the course of 7 days (N=12 Taste buds) along with standard deviation.

### Intravital imaging reveals changes in the morphology and volume of gustatory fiber innervation after CYP treatment

To determine how gustatory nerve fibers changed over the course of CYP treatment, six Phox2b-Ai9 mice were imaged to identify trackable taste buds. These mice were then separated into two groups, with the treatment group injected with CYP and the control group injected with saline. Z-stacks of tracked taste buds were imaged 2- and 4-days post treatment (Figure 5A). The gustatory fibers of saline-injected mice are consistent in their volume over the course of imaging (Figure 5B). The CYP-treated group showed a significant decrease in the volume of gustatory fibers within individual taste buds 4-days post treatment (Figures 5C). Figure 5D shows the percent change of Phox2b-Ai9 fiber volume in individual taste buds over time for both CYP-treated and control mice. To further examine the CYP-induced fiber loss, one saline- and one CYP-treated mouse were selected. Taste maps were taken to track the individual taste buds over treatment (Figure 5E, G). As seen in our previous results, fiber volume remained consistent after multiple days of imaging in saline injected mice (Figure 5F). CYP treated taste buds showed variability in how they were affected by the treatment, with some of the taste bud showing consistent fiber volume and morphology even after treatment (Supplemental Figure 1), and others showing a significant decrease in their volume (Figure 5H). These results support the hypothesis that gustatory fibers decrease after cyclophosphamide treatment.

**Figure 5:**
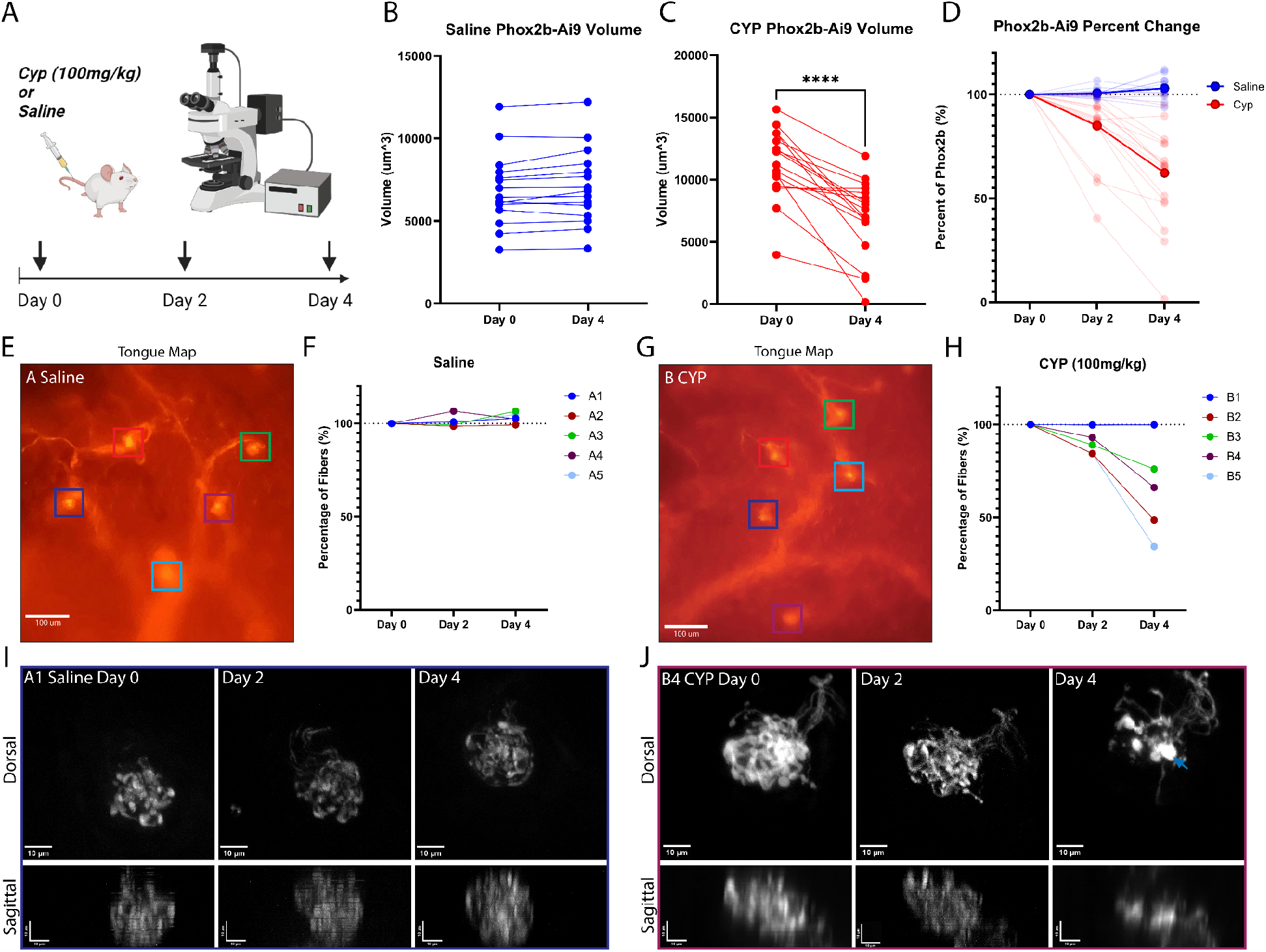
Intravital Imaging Reveals Decreased Volume of Gustatory Fiber Innervation after CYP Treatment. **A**. 6 mice were separated into two groups, CYP- and saline-treated. Taste buds were then selected and imaged for day 0. The mice were then injected with either CYP (100 mg/kg) or saline and imaged 2- and 4-days post-injection. **B, C**. The volume of taste buds for mice injected with CYP and saline, each line representing one taste bud (N=15-17 TBs per group). Wilcoxon ranked test were used to show significance (*p<0.05). **D**. The percentage change per taste bud for saline vs CYP taste buds over a 4 day period. Solid line showing the averages, transparent lines showing individual taste buds. **E**. Tongue map from a saline-injected mouse (Mouse A), showing 5 tracked taste buds in different colored boxes. **F**. The percent change per taste bud for saline-treated mouse over 4 days **G**. Tongue map from a CYP-injected mouse (Mouse B), showing 5 tracked taste buds in different colored boxes. **H**. The percent change per taste bud for CYP-treated mouse over 4 days **I**. a1 (blue) taste bud from the saline-treated mouse tracked for 4 days, showing maximum projection of the z-stack for dorsal, re-sliced maximum projection of sagittal images. **J**. b4 (purple) taste bud from the Cyp-treated mouse tracked for 4 days, showing maximum projection of the z-stack for dorsal, re-sliced maximum projection of sagittal images. Large retraction bulbs within the taste bud’s neuronal fiber core (blue arrow).

After examining the morphology of treated taste buds, saline treated taste buds remain consistent over the course of imaging, as seen in our previous results (Figure 5I). CYP-treated taste buds were variable in their morphology response, where one taste bud showed a minor decrease 2 days post-treatment. However, by day 4, there was a major loss in the gustatory fibers within the taste bud, showing a loss in z-height (Figure 5J). Large retraction bulbs are shown to appear within the core of the taste bud (blue Arrow). Next, tracked taste buds were taken and whole-mount stained with TROMA as previously described. Saline-tracked taste buds showed consistent gustatory fiber volume from the 2-photon imaging (Figure 6A). These taste buds remained intact, with the gustatory fibers innervating the TRCs (Figure 6B). One CYP-treated taste bud showed a decrease in volume by 4 days post treatment, with the fibers spreading out from the fiber core, showing an increased diameter of the taste bud’s fiber reach (Figure 6C). Examining the whole-mount staining of this taste bud reveals the taste bud to have fractured, with the gustatory fibers appearing to be extending from the core to follow the remaining TRCs (Figure 6D). All the results together show a significant decrease in gustatory fiber volume after CYP treatment, although it is variable to how severe this loss is dependent on the taste bud.

**Figure 6:**
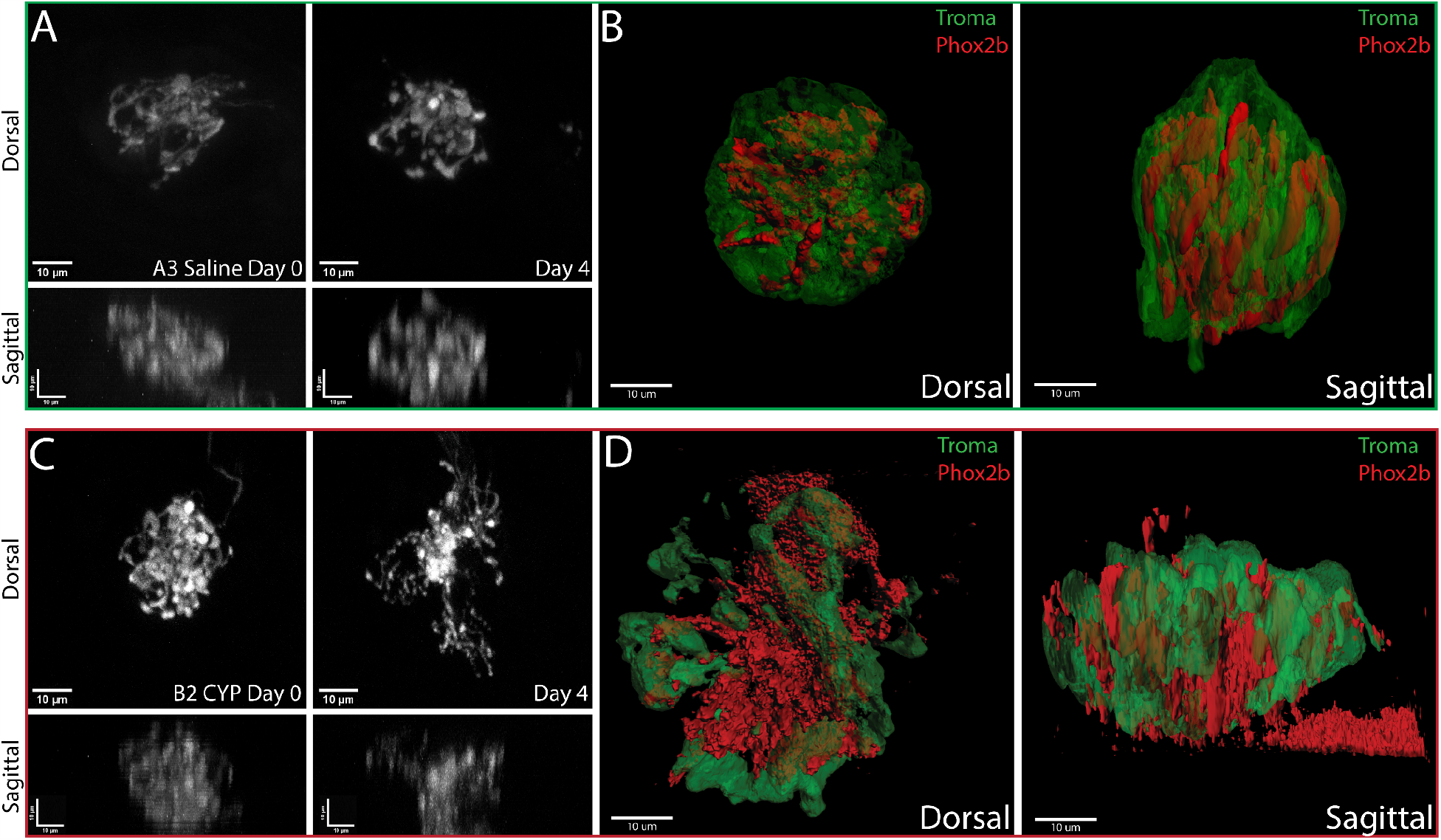
Morphology Changes Within CYP-Treated Taste Buds Seen in TRCs and Gustatory Fibers. **A**. A3 (green) taste bud from a saline-treated mouse, showing maximum projection of the z-stack for dorsal, re-sliced maximum projection of sagittal images for 0- and 4-days **B**. 3D-reconstruction of the taste bud 4 days after saline injection, TRCs shown in green, and gustatory fibers shown in red, showing dorsal and sagittal sides of the taste bud. **C**. B2 (red) taste bud from CYP-treated mouse, showing maximum projection of the z-stack for dorsal, re-sliced maximum projection of sagittal images for 0 and 4 days. **D**. 3D-reconstruction of the taste bud 4 days after CYP injection, TRCs shown in green, and gustatory fibers shown in Red, showing dorsal and sagittal sides of the taste bud.

## Discussion

Consistent with previous reports, we found that a single dose of 100mg/kg cyclophosphamide produces a loss (within 4-8 days) and gradual repopulation (within ∼20 days) of Type II and Type III taste receptor cells in the fungiform and circumvallate papillae^7-9,20^. Interestingly, we found that peripheral nerve fibers are reduced within the taste bud proper at 4 and 8 days post treatment, but are re-established by 20 days post-treatment. Intravital imaging of Phox2b-Cre;tdTomato labeled gustatory fibers innervating the fungiform taste buds revealed that some buds were more affected than others, but that fibers changed their morphology in a way consistent with what had been previously observed with sonidegib treatment – a retraction and splaying out and spreading of fibers at the core/base of the epithelium^15^.

These results suggest that gustatory fibers are indeed sensitive to chemotherapy treatment. Gustatory fiber volume is reduced within the taste bud proper by 4 days of treatment. This coincides, or even precedes, the loss of taste receptor cells within the bud. While the average reduction in fiber innervation is significant when averaged across animals, individual buds vary in the degree of severity. This variability was more obvious when observed in whole-mount preparations of the fungiform taste papillae and using 2-photon intravital imaging.

To our knowledge, this is the first report detailing the effects of CYP on gustatory nerve fibers within taste buds. The closest comparable studies have used another agent, sonidegib, which is a hedgehog pathway inhibitor and is used to treat some basal cell carcinomas. Long-term treatment (25-28 days) of sonidegib (oral gavage 20mg/kg) inhibits the maturation of taste receptor cells, and eventually eliminates the taste bud entirely. Although the chorda tympani nerve in sonidegib treated rodents is no longer sensitive to chemical (taste) stimuli, mechanosensory and thermosensory responses are maintained^21-23^. After sonidegib treatment, the morphology of phox2b-Cre;TdTomato labeled chorda tympani fibers change dramatically, reorganizing and expanding just under the fungiform papilla epithelium where taste buds were ^15^. Our study uses a single dose of CYP, which has a less dramatic effect on the TRCs within the bud compared to long-term sonidegib. Nevertheless, we observed several examples of taste buds where the fibers also flatten and spread out at the basal side of the bud. In our experiments, this reorganization happens within 4 days of CYP treatment. In the sonidegib study, increased expression of GAP43 indicated that these fibers were in the process of remodeling. It will be interesting to test the time course of GAP43 expression in the fibers after CYP treatment during this period as well, or later in the recovery phase 8-20 days post CYP.

Another recent study used 2-photon intravital imaging to track the dynamics of individual gustatory fibers in the taste buds^24^. Similar to Whiddon et al, we also find that the terminal fibers within the taste bud are surprisingly dynamic. While our study does not attempt to track the dynamics of individual fibers innervating the bud as Whiddon does, we do see a remarkable amount of motility and change of the fibers within the bud, while the overall volume of innervation is maintained (at least for the saline treated control mice). This study also used sonidegib (LDE225) treatment to perturb the taste buds. Interestingly, 10 days after LDE225 treatment, no major differences in the rate of terminal branch changes were reported compared to controls. And even 20-40 days after treatment, the dynamics of terminal arbors remained regardless of whether the taste bud had recovered or not. These data are somewhat inconsistent with Donnelly et al, but this may be due to a shorter sonidegib treatment window (10 days vs 25 days) or the fact that only individual nerve fibers were labeled, rather than the entire Phox2b+ population^15^. Together, these studies highlight the usefulness and promise of intravital imaging to investigate important questions about the dynamics of taste bud remodeling in normal animals and in those where the taste system is perturbed.

Loss of taste function is also commonly reported by patients undergoing radiotherapy treatment for head and neck cancers. Fractionated and single-dose irradiation in mice reduces taste progenitor cell numbers and their proliferation, interrupting the supply of new taste receptor cells to taste buds^25,26^. These studies did not investigate the effects of radiotherapy on the gustatory fibers within taste buds, but it will be important to see if both radio- and chemotherapies have similar or different effects. This will have important implications for future studies investigating the mechanisms by which gustatory fibers are lost/retracted from taste buds during cancer treatment.

Gaillard et al suggest a role for Wnt/B-catenin and potentially other signaling pathways to maintain taste stem cell proliferation during radiation exposure and/or promote taste cell differentiation following radiation injury^25^. Importantly, R-spondin is expressed and secreted by gustatory neurons and potentiates the Wnt pathway^27,28^. R-spondin has also been shown to be the neuronal factor necessary for taste bud homeostasis^27,28^. Interestingly, treatment with Rspo1 is protective for chemotherapy- and radiotherapy-induced oral mucositis^29^. This points to a potentially important role of gustatory neurons to produce R-spondin and promote the differentiation of taste stem cell populations to recover taste buds after insults like chemo and radiotherapy. It will be important to determine if the remodeling/migration of gustatory fibers toward the base of the taste bud after CYP treatment that we observed serves a restorative function, stimulating the remaining stem cell populations to generate more mature TRCs and lingual epithelial cells.

Another important factor that may influence the loss and recovery of taste function after chemotherapy treatment is inflammation. Cyclophosphamide treatment in mice quickly induces TNF-α expression, especially by Type II TRCs. TNF-α levels peak at 8-24 hours post CYP injection, but persist at elevated levels for ∼72 hours^30^. Pre-treatment with amifostine, a sulfhydryl drug that protects tissues during chemo- or radiation therapy, reduced the amount of CYP-induced TNF-α expression in taste buds^30^, and has been shown to improve taste function in CYP-treated mice^7^. It is very likely that inflammation may also reduce the ability of gustatory neurons to properly innervate the TRCs within the taste buds or otherwise disrupt taste signaling, just as inflammation affects other peripheral sensory neurons such as nociceptors^31-35^. It will be important to understand the relationship between inflammation and gustatory neuron innervation both in the context of chemotherapy/radiotherapy treatment, but also for other inflammatory conditions that can affect the taste system including obesity, diabetes, and metabolic syndrome, among others.

In summary, we find that gustatory nerve fibers innervating taste buds are sensitive to cyclophosphamide treatment, reducing innervation of the taste buds within 4-8 days, but re-innervating the buds at normal levels by 20 days post treatment. Individual taste buds in the fungiform papilla differ in their response to CYP, with some maintaining near-normal levels of innervation, and others losing the majority of their innervation within the taste bud proper. Future experiments will be needed to determine the mechanisms and sequence of events leading to fiber retraction as it relates to taste receptor cell loss within the buds.

Fibers may begin to retract earlier than TRCs die, but perhaps subsequent to inflammatory signaling events. To further investigate if and how the taste buds fully regain function, it will be necessary to investigate the period between 8- and 20-days post treatment more carefully, especially given that many chemotherapy patients report prolonged taste dysfunction long after their treatment concludes. It will also be important to determine if this effect is generalizable across multiple types of chemotherapy agents and radiation-based treatments for cancers.

## Figures

**Supplemental figure 1:**
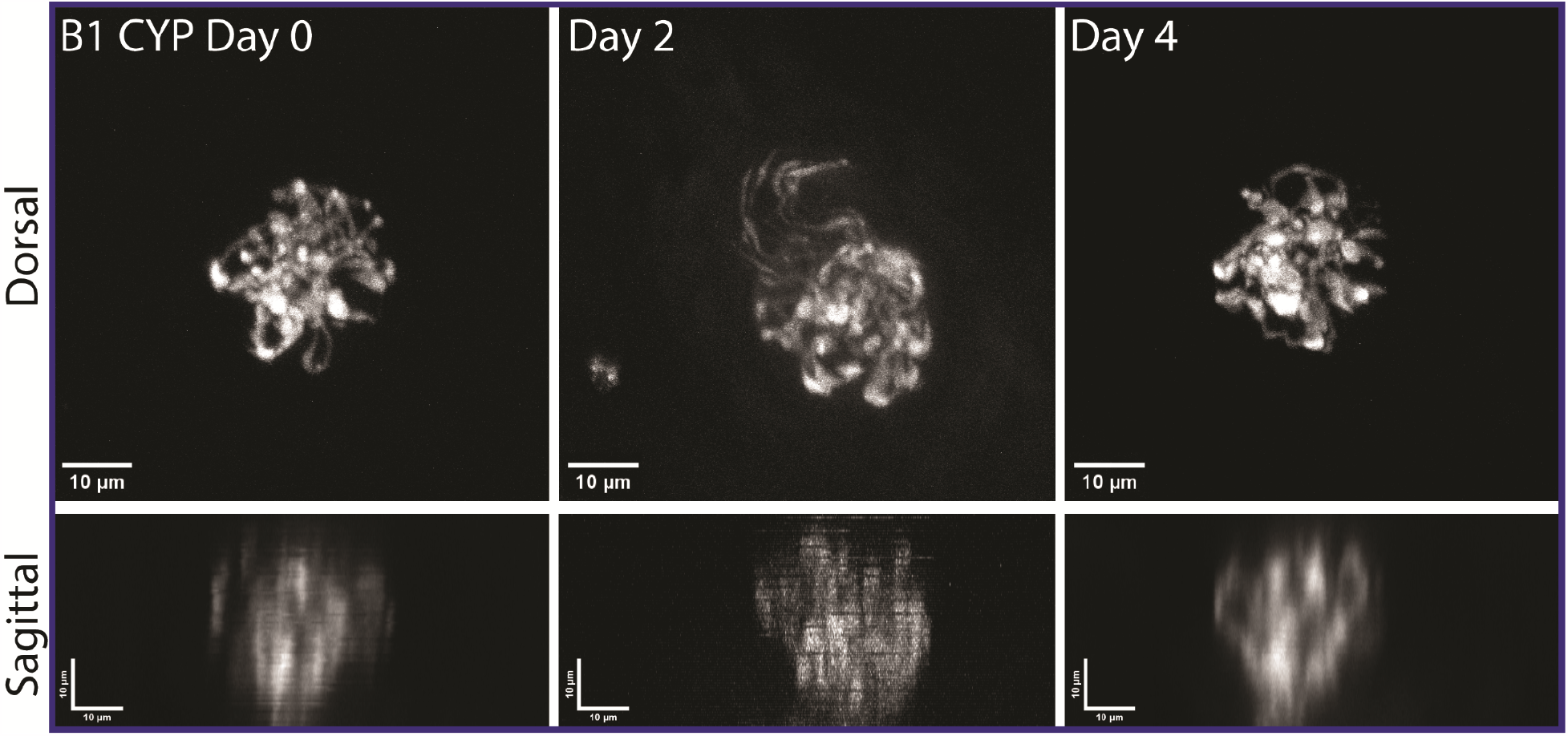
Variability Between CYP-Treated Taste Buds Seen in Gustatory Fibers ^i^B1 (Blue) taste bud from CYP-treated mouse on days 0, 2, and 4, showing maximum projection of the z-stack for the dorsal view and re-sliced maximum projection for sagittal views.

